# Population genetic structure in the insular Ryukyu flying fox, *Pteropus dasymallus*

**DOI:** 10.1101/2020.03.05.979211

**Authors:** Shiang-Fan Chen, Chung-Hao Juan, Stephen Rossiter, Teruo Kinjo, Dai Fukui, Kuniko Kawai, Susan M. Tsang, Maria Josefa Veluz, Hiroko Sakurai, Nian-Hong Jang-Liaw, Keiko Osawa, Wen-Ya Ko, Masako Izawa

## Abstract

Small isolated populations are vulnerable to both stochastic events and the negative consequences of genetic drift. For threatened species, the genetic management of such populations has therefore become a crucial aspect of conservation. Flying foxes (*Pteropus* spp, Chiroptera) are keystone species with essential roles in pollination and seed dispersal in tropical and subtropical ecosystems. Yet many flying fox species are also of conservation concern, having experienced dramatic population declines driven by habitat loss and hunting. The Ryukyu flying fox (*Pteropus dasymallus*) ranges from Japan and Taiwan to the northern Philippines, and has undergone precipitous population crashes on several islands in recent decades. To assess population genetic structure and diversity in *P. dasymallus*, and its likely causes, we analyzed mitochondrial and microsatellite DNA. Both markers showed significant genetic differentiation among most island populations with patterns of isolation-by-distance. However, while mitochondrial haplotypes showed some mixing across the region, likely reflecting historical colonization and/or dispersal events, microsatellites markers showed clear subdivisions corresponding to the position of deep ocean trenches. The current distribution of *P. dasymallus* and its subspecific diversity therefore appears to have arisen through vicariance coupled with a long history of restricted gene flow across oceanic barriers. We conclude that isolated island subgroups should be managed separately, with efforts directed at reducing further declines.

## Introduction

Small and isolated populations are vulnerable to stochastic events and the effects of genetic drift, potentially leading to the loss of diversity, reduced reproductive fitness, and increased risks of extinction (Ellstrand & Elam 1993; Frankham 2010; Jordan et al. 2016). The genetic management of such populations is therefore considered an essential aspect of conservation, particularly for endangered and threatened species. With detailed genetic information, we can effectively monitor the loss of genetic diversity and precisely estimate population parameters such as population size fluctuation, admixture, and gene flow, all of which contribute to our understanding of long-term species survival (Allendorf et al. 2010; Shafer et al. 2015). Genetic and nongenetic (e.g., behavioral, ecological, demographic, and environmental) considerations can therefore be integrated to enhance the efficiency of conservation programmes and further formulate appropriate management strategies (Hoban et al. 2013; Polechová & Barton 2015; Frankham et al. 2017).

Old World fruit bats (Chiroptera: Pteropodidae) are keystone species that play essential roles in pollination and seed dispersal in tropical and subtropical ecosystems (Cox et al. 1991; Fujita & Tuttle 1991). Aside from promoting long-distance seed dispersal needed for forest restoration (Nyhagen et al. 2005; Shilton & Whittaker 2009), these species also pollinate a number of economically important plants, including durian (Aziz et al. 2017). Yet despite acting as major providers of ecosystem services, Old World fruit bats face a range of threats. Over recent decades, the combined impacts of habitat loss, forest degradation, and hunting for bushmeat have all led to severe and rapid population declines (Mickleburgh et al. 2002), with wide-ranging negative ecological and other impacts (Cox & Elmqvist 2000; McConkey & Drake 2006; Florens et al. 2017). Currently, of the nearly 200 recognised species, over a half are of conservation concern, including around 30 *Pteropus* species (IUCN 2019).

Larger-bodied Old World fruit bats are generally considered to be strong and capable fliers with extensive home-ranges. For example, genetic analyses of *P. scapulatus* and *Eidolon helvum* have suggested gene flow can occur over thousands of kilometres, across mainland Australia and Africa, respectively (Sinclair et al. 1996; Peel et al. 2013). Although long distance movements in large Old World fruit bats might result from natal dispersal events, or from storms and typhoons, their capacity for long distance day-to-day movements, even among islands, is likely to be an adaptive trait for tracking ephemeral food resources. For example, long-distance inter-island movements have been recorded and/or inferred indirectly from genetic data in several *Pteropus* species, e.g., *P. dasymallus inopinatus, P. medius, P. niger, P. tonganus*, and *P. vampyrus* (McConkey & Drake 2007; Nakamoto et al. 2011a; Larsen et al. 2014; Tsang et al. 2018; Olival et al. 2019). In these taxa, populations on adjacent islands can appear as single panmictic units. However, in other flying fox species, e.g., *P. livingstonii, P. mariannus*, and *P. samoensis*, genetic structure can be present among island groups, indicative of restricted gene flow (Brown et al. 2011; Russell et al. 2016; Ibouroi et al. 2018). These contrasting scenarios require different conservation management approaches; in the former case, island groups might be managed as a single entity, while in the latter case, islands populations might be better treated as distinct evolutionarily significant units (ESU), and thus managed separately (Epstein et al. 2009; Oleksy et al. 2019).

The Ryukyu flying fox (*Pteropus dasymallus*) is distributed from the Ryukyu Archipelago of Japan through Taiwan to the northern Philippines (Kinjo & Nakamoto 2009; Figure 1). Five subspecies are recognized, with populations from Ryukyu Archipelago classified into four subspecies based on their respective island group ranges (Daito flying fox, *P. d. daitoensis*; Erabu flying fox, *P. d. dasymallus*; Orii’s flying fox, *P. d. inopinatus*; and Yaeyama flying fox, *P. d. yayeyamae*). The fifth subspecies, from Taiwan, is recognized as the Formosan flying fox (*P. d. formosus*) (Yoshiyuki 1989; Mickleburgh et al. 1992), while a population in the Philippines has been discovered more recently and has yet to be named formally as a subspecies (Heaney et al. 1998). This latter population occurs north of Luzon on two oceanic island groups, the Batanes and Babuyan Islands, and its unclear subspecific status in part reflects the logistical difficulty in traveling to these islands.

**Figure 1.**
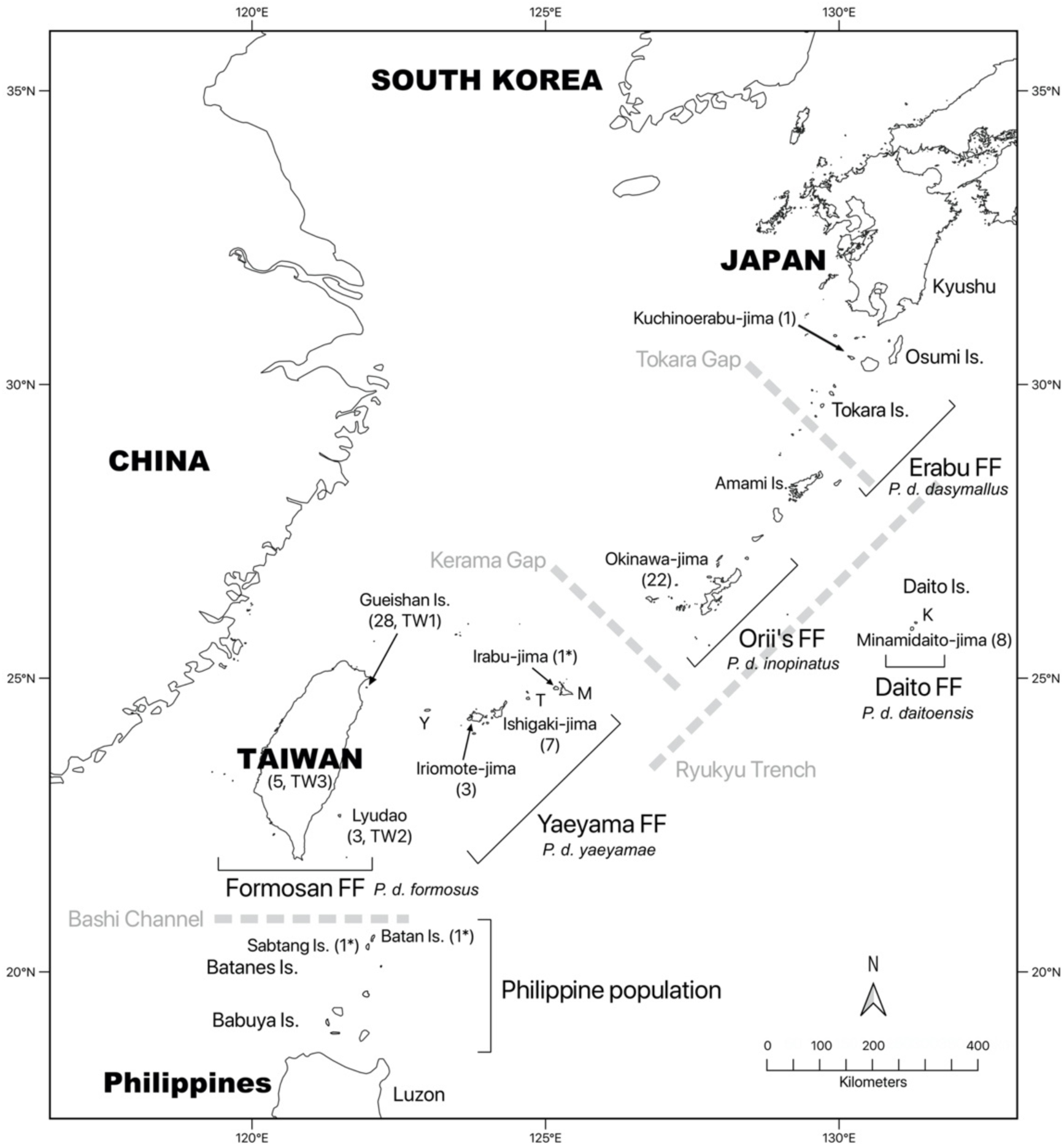
Map of the Ryukyu Archipelago, Taiwan, and northern islands of the Philippines showing the distribution of *Pteropus dasymallus*. The locations where the samples used in this study originated is also shown with corresponding sample size presented in brackets. An asterisk indicates DNA sequences acquired from Genbank or Dryad. Abbreviation: FF, flying fox; Y, Yonaguni-jima; T, Tarama-jima; M, Miyako-jima; K, Kitadaito-jima.

*P. dasymallus* is currently categorized as Vulnerable by the IUCN Red List (IUCN 2019), however, the local conservation status differs among the five subspecies (Vincenot et al. 2017). In particular, the Daito, Erabu, and Formosan flying foxes are all characterized by small populations numbering approximately 100-300 individuals (Saitoh et al. 2015). All three are also protected by national laws, with the Daito and Erabu flying foxes designated as ‘Natural Monuments’ of Japan, and the former also designated as a ‘National Endangered Species’. Similarly, the Formosan flying fox is also afforded protection, designated as an ‘Endangered Species’ in Taiwan. In contrast, the two subspecies Orii’s and Yaeyama flying foxes, and the Philippine population, appear more common and are not classified as locally threatened (Heaney et al. 1998; Nakamoto et al. 2011b; Saitoh et al. 2015).

The Daito flying fox is geographically restricted to only two small islands of the Daito Islands (Minamidaito-jima and Kitadaito-jima, Figure 1), where most natural habitat has been converted into farmland, and where typhoons are the major current threat (Saitoh et al. 2015). The Erabu flying fox is found on the Ōsumi Islands and Tokara Islands, representing the northern limit of this species (Yoshiyuki 1989). In Taiwan, the Formosan flying fox was once abundant in its original main habitat on Lyudao (Green Island) 30.6 km off the southeastern coast of Taiwan; however, this island population experienced dramatic hunting and habitat loss in the 1970s and 1980s (Lin & Pei 1999), leading to its near extinction, with only four individuals have been recorded in recent years (Chen 2009). In 2004, an additional small population of the Formosan flying fox was recorded for the first time on Gueishan Island (Turtle Island), 9.7 km off the northeastern coast of Taiwan, and some individuals have occasionally been found on the main island of Taiwan since 2006 (Wu 2010). At the present time, nothing is known about the origin of the flying foxes on Gueishan Island and Taiwan’s main island.

To date, the phylogenetic relationships and population divergence among the subspecies of *P. dasymallus* remains obscure but is likely to reflect the geography of the Ryukyu Archipelago. This island chain comprises around 150 islands, extending for 1200 km between the main islands of Japan to Taiwan. Although several adjacent Ryukyu islands were connected by a land bridge in the Last Glacial Maximum (LGM) (Ota 1998), other islands remained isolated from each other. In fact, genetic studies of several taxonomic groups have revealed divergence between populations from the Northern, Central, and Southern Ryukyus (Ota 1998, 2000), areas which remained separated in the LGM by two deep tectonic straits, the Tokara Gap (Tokara Strait) and the Kerama Gap (Miyako Strait) (Figure 1). The Tokara Gap lies between Akuseki-jima and Kodakara-jima in the Tokara Islands, while the Kerama Gap, the widest strait in the Ryukyu Islands, lies between Okinawa-jima and Miyako-jima (Nakamura et al. 2013). Therefore, both of these gaps are likely to have formed geographical barriers, so promoting genetic drift and divergence among the Northern, Central and/or Southern Ryukyu populations (Toda et al. 1997; Lin et al. 2002). Of these, Taiwan is geographically closest to the Southern Ryukyus (110 km), and both species and genetic diversity have been found to be more similar among Taiwan and the Southern Ryukyus than between these islands and other regions (>270 km) (Ota 2000; Tominaga et al. 2015). To the south, the two oceanic island groups of the Batanes and the Babuyan Islands were formed about three million years ago (late Pliocene), with continued uplift and volcanism into the Pleistocene. The Batanes is closer to the southern tip of Taiwan (150 km, Bashi Channel) than they are to Luzon (200 km) (Bellwood & Dizon 2013). However, the genetic structure of mammals on the island chain has seldom been addressed (Yoshikawa et al. 2016).

Here we examined genetic diversity and population genetic structure in *P. dasymallus* using mitochondrial DNA (mtDNA) and microsatellite markers. We aimed to (1) assess the genetic diversity of *P. dasymallus* across the different island groups, (2) examine whether genetic differentiation exists among these groups in line with their subspecies designations, (3) examine the pattern of genetic structure, and (4) examine the relationships between both the newly-recorded Formosan flying fox individuals (from Gueishan Island and Taiwan’s main island) and the little-studied Philippine samples, and the other subspecies. Our hypotheses were as follows: First, we hypothesized that the Daito, Erabu, and Formosan flying fox subspecies will show lower genetic diversity than the other subspecies due to their relatively small population sizes. Second, we hypothesized that genetic differentiation exists in *P. dasymallus* and corresponds to the respective subspecies identities. Third, like in many other species in this region, this differentiation can be also accounted for by regional deep sea trenches: the Tokara and Kerama Gaps in the Ryukyu Archipelago, and the Bashi Channel between Taiwan and the northern Philippines. Finally, we hypothesized that the Formosan flying fox individuals, newly recorded on Gueishan Island and Taiwan’s main island, will share a similar genetic structure with bats from the nearest Yaeyama Islands, suggesting a colonization event. An understanding of the genetic structure and degree of gene flow among different island groups of *P. dasymallus*, along with insights into whether these patterns are ancient or new, can help to inform conservation management decisions, including whether or not to treat small populations separately, or whether to translocate isolated vagrants.

## Methods

### Sampling

We obtained samples of *P. dasymallus* opportunistically over a period of 10 years (2009-2019) from wild-caught individuals and carcasses found in the wild, as well as rescued and/or captive individuals. All samples originated from eight different Taiwanese and Ryukyu islands, and were classified into the five subspecies based on their geographical source following Mickleburgh et al. (1992). A total of 77 samples were analyzed for this study after removing duplicate samples and putative parents or offspring of other individuals, as outlined below. Sample sizes per subspecies were 36 Formosan, 10 Yaeyama, 22 Orii’s, 1 Erabu, and 8 Daito. Formosan flying fox samples were further divided into three groups denoted as TW1 (Gueishan Island), TW2 (Lyudao), and TW3 (Taiwan’s main island) according to the islands from which they originated. Yaeyama flying fox samples originated from Iriomote-jima and Ishigaki-jima, Orii’s flying fox samples from Okinawa-jima, Erabu flying fox samples from Kuchinoerabu-jima, and Daito flying fox samples from Minamidaito-jima (Figure 1, Appendix 1).

Samples ranged from wing membrane biopsies, blood and frozen muscle tissue to fecal samples. Wing membrane samples were collected with a 3-mm biopsy punch and placed in 99.5% ethanol, Allprotect Tissue Reagent (Qiagen), or silica beads until extraction. For the blood samples, a volume of 0.5 cc was taken by a professional veterinarian and preserved in ethylenediaminetetraacetic acid (EDTA) anticoagulant. Frozen muscle tissue obtained from specimens, and fresh feces, were stored in 99.5% ethanol or RNAlater RNA Stabilization Reagent (Qiagen).

### DNA extraction and amplification

To extract genomic DNA from wing membrane, blood, and frozen muscle samples, we used DNeasy Blood and Tissue Kits (Qiagen). For fecal samples, we used the QIAamp Investigator Kit or QIAamp Fast DNA Stool Mini Kit (Qiagen).

We amplified a section of the mtDNA control region using the primers BovL 14987 (5’-CGC-ATA-TGC-AAT-CCT-ACG-A-3’) and BovR 15967 (5’-GCG-GGT-TGC-TGG-TTT-CAC-3’), which we designed for this study. Polymerase chain reaction (PCR) was carried out in a total volume of 15 μl, containing 20-100 ng of template DNA, 0.25μl of 10 μM of each primer, and 7.5 ul of Quick Taq HS DyeMix (TOYOBO). Amplification was performed with the following profile: 2 min at 94ºC followed by 30 cycles of 30 s at 94ºC, 30s at annealing temperature (55ºC), 50 s at 68ºC, and a final extension of 10 min at 68ºC. PCR products were run on an ABI 3730XL DNA Analyzer (Applied Biosystems). The chromatograms were edited and aligned in the program of SeqMan and MegAlign (DNASTAR). We also obtained three published *P. dasymallus* partial control region sequences from GenBank and Dryad, for one Yaeyama flying fox from Irabu-jima (accession NC_002612.1) (Nikaido et al. 2000), and two individuals collected from the Batanes Islands, with one from Batan Island and the other from Sabtang Island (accessions MJV458 and MJV451, respectively), representing the Philippine population (Tsang et al. 2019).

For microsatellite DNA analysis, 108 species-specific markers were generated by Genetic Identification Services (CA, USA). To quantify polymorphism and characteristics of these loci, we used a subset of samples. In total, 26 loci were polymorphic and used for subsequent genotyping 76 samples (Supporting Information S1). For this, PCRs were carried out in a total volume of 10 μl, containing approximately 10-50 ng of template DNA, 0.5 μl of 10 μM of each primer, and 5 μl of Quick Taq HS DyeMix. Amplification was performed with the following profile: 2 min at 94°C, followed by 40 cycles of 30 s at 94°C, 30 s at annealing temperature (54°C), 1 min at 68°C, and a final extension of 10 min at 68°C. PCR products were also run on an ABI 3730XL DNA Analyzer, and allele scoring was performed using the software GeneMarker 4.2 (SoftGenetics). Identity and parentage analyses were performed using Cervus 3.0.7 (Kalinowski et al. 2007) to identify duplicate samples and parentage. Samples with exactly matching genotypes across all loci were determined as duplicates, and removed. Parentage was determined based on no allele mismatches. Only one sample from each duplicate or parent-offspring pair was included for further analyses.

### MtDNA analysis

Based on mtDNA data, we estimated the number of haplotypes, haplotype diversity (*h*), nucleotide diversity (π), and average number of pairwise differences for each subspecies. We determined the extent of genetic differentiation by applying an analysis of molecular variance (AMOVA) (Excoffier et al. 1992). Only populations with sample sizes greater than one were included. The total variance was partitioned into variance components attributable to within and among subspecies. To measure the degree of genetic differentiation among subspecies, the derived index of the total population was estimated. The significance of the differentiation was tested by performing 20,000 random permutations. These analyses were performed using Arlequin 3.5.2.2 (Excoffier & Lischer 2010).

We also estimated pairwise differentiation among subspecies, and examined isolation by distance for genetic distances, estimated by Φ_ST_, among islands. For geographical distances (km) among pairwise islands, linear Euclidean distances between the centers of pairwise sampling islands were computed based on latitudinal and longitudinal coordinates. A total of eleven islands were included in the analysis of isolation by distance (Gueishan Island, Lyudao, the main island of Taiwan, Iriomote-jima, Ishigaki-jima, Irabu-jima, Okinawa-jima, Kuchinoerabu-jima, Minamidaito-jima, Batan Island, and Sabtang Island). The significance level was assessed using a Mantel test with 20,000 permutations in Genepop (web version) (Raymond & Rousset 1995; Rousset 2008).

To further visualize genetic structure with respect to subspecies, we generated a haplotype network. An unrooted maximum likelihood tree was generated in MEGA X (Kumar et al. 2018) and converted into a haplotype network using Haplotype Viewer (Center for Integrative Bioinformatics Vienna).

### Microsatellite DNA analysis

Deviation from Hardy-Weinberg equilibrium (HWE) at each microsatellite locus and subspecies, and linkage disequilibrium for each pair of loci, were tested using the Markov chain method (10,000 dememorization steps, 1,000 batches and 10,000 iterations per batch). We assessed statistical significance using Bonferroni correction for multiple comparisons. For each microsatellite locus, we recorded the number of alleles (N_A_), observed heterozygosity (H_O_), and expected heterozygosity (H_E_). For each subspecies, we derived diversity indexes, including the mean number of alleles (N_a_), allelic richness corrected for unequal sample size (A_C_), and the mean H_O_ and H_E_. The average pairwise relatedness (RI) of each subspecies was calculated to infer relationships between individuals (Ritland 1996).

Like mtDNA, we also conducted genetic structure analyses, including AMOVA, pairwise differentiation, estimated by F_ST_, and isolation by distance, for microsatellite data. Eight islands were included here (Irabu-jima, Batan Island, and Sabtang Island were excluded, where no microsatellite data was available). These analyses were performed in GenAlEx 6.51 (Peakall & Smouse 2006, 2012) or Genepop (web version).

To examine relationships among populations based on multilocus microsatellite genotype data, we inferred the number of genetically distinct clusters using the Bayesian clustering approach implemented in STRUCTURE 2.3.4 (Pritchard et al. 2000; Falush et al. 2003). An admixture ancestry model with correlated allele frequencies was used with a burnin period of 100,000 iterations followed by 1,000,000 Markov chain Monte Carlo (MCMC) repetitions. The number of ancestral populations (*K*) was set to 1 to 10. Ten independent runs for each *K* to confirm consistency across runs were performed with prior information on population origins. The best number of *K* was determined based on the mean likelihood (*L*(*K*)) and variance for each *K* value and the *ad hoc* statistic Δ*K* with the Evanno method using the program Structure Harvester (Evanno et al. 2005; Earl & vonHoldt 2012). The output data were generated and visualized with Clumpak 1.1 (Kopelman et al. 2015).

## Results

### MtDNA analysis

Analyses of partial mtDNA control region sequences revealed 33 haplotypes with 22 parsimony informative sites from a total of 80 *P. dasymallus* samples encompassing the five recognized subspecies and the Philippines population. Haplotype diversity (*h*), nucleotide diversity (π), and pairwise difference averaged over all samples were 0.948, 0.012, and 3.556, respectively. A summary of the genetic diversity is presented in Table 1. The Philippine population showed the highest diversity. On the other hand, the Daito flying fox consistently showed the lowest diversity. We excluded the Erabu flying fox individuals from the subspecies-level analyses given that only one sample is available.

**Table 1.**
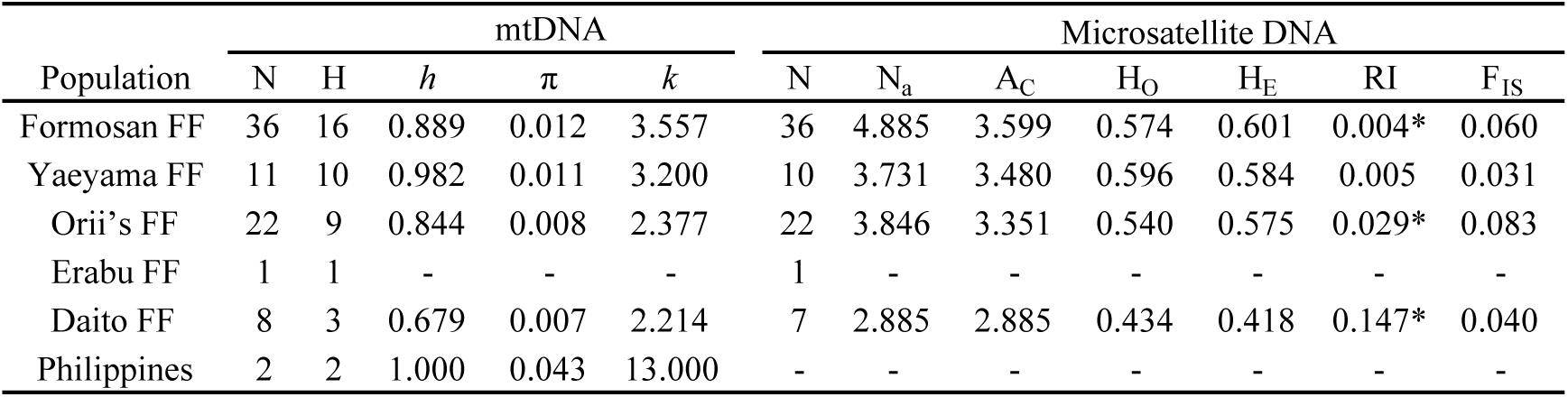
Genetic diversity of *Pteropus dasymallus* subspecies estimated with mitochondrial DNA control region and 26 microsatellite loci. N: sample size, H: number of haplotype observed, *h*: haplotype diversity, π: nucleotide diversity, *k*: average number of nucleotide differences, N_a_: mean number of alleles per locus, A_C_: allelic richness, H_O_: observed heterozygosity, H_E_: expected heterozygosity, RI: relatedness, F_IS_: inbreeding coefficient. The Erabu flying fox was excluded for subspecies-level analyses given that only one sample is available. Abbreviation: FF, flying fox.

| Population | mtDNA |  |  |  |  | Microsatellite DNA |  |  |  |  |  |  |
| --- | --- | --- | --- | --- | --- | --- | --- | --- | --- | --- | --- | --- |
| | N | H | $h$ | $\pi$ | $k$ | N | $N_a$ | $A_C$ | $H_O$ | $H_E$ | RI | $F_{IS}$ |
| Formosan FF | 36 | 16 | 0.889 | 0.012 | 3.557 | 36 | 4.885 | 3.599 | 0.574 | 0.601 | 0.004* | 0.060 |
| Yaeyama FF | 11 | 10 | 0.982 | 0.011 | 3.200 | 10 | 3.731 | 3.480 | 0.596 | 0.584 | 0.005 | 0.031 |
| Orii's FF | 22 | 9 | 0.844 | 0.008 | 2.377 | 22 | 3.846 | 3.351 | 0.540 | 0.575 | 0.029* | 0.083 |
| Erabu FF | 1 | 1 | - | - | - | 1 | - | - | - | - | - | - |
| Daito FF | 8 | 3 | 0.679 | 0.007 | 2.214 | 7 | 2.885 | 2.885 | 0.434 | 0.418 | 0.147* | 0.040 |
| Philippines | 2 | 2 | 1.000 | 0.043 | 13.000 | - | - | - | - | - | - | - |

The AMOVA revealed genetic differentiation among the five analyzed populations (the Formosan, Yaeyama, Orii’s, and Daito flying foxes and Philippine population with sample sizes greater than one). The Φ_ST_ value was 0.140, which was significantly different from zero (P < 0.001, Table 2). This indicated that approximately 14.0% of the total mtDNA genetic variation was accounted for by the differences among subspecies. The magnitude of the pairwise differentiation varied markedly with the lowest value shown in the pair of the Formosan and Yaeyama flying foxes. The Daito flying fox and Philippine population presented relatively high values of differentiation with other counterparts (Table 3).

**Table 2.** Analysis of molecular variance (AMOVA) for *Pteropus dasymallus*. Five populations, including the Formosan, Yaeyama, Orii’s, and Daito flying foxes and Philippine population, with sample sizes greater than one are included in the mtDNA analysis. The first four are included in the microsatellite analysis.

| Source of | mtDNA |  | Microsatellite DNA |  |
| --- | --- | --- | --- | --- |
|  | variation | P value | variation (%) | P value |
| Among | 13.97 | < 0.001* | 6.91 | < 0.001* |
| Within subspecies | 86.03 |  | 93.09 |  |

\*: statistically significant

**Table 3.** Pairwise genetic differentiation between *Pteropus dasymallus* subspecies or population. Φ_ST_, above diagonal based on mtDNA data; F_ST_, below diagonal based on microsatellite data. The Erabu flying fox is excluded. Statistical significance is also provided. Abbreviation: FF, flying fox.

| Population | Formosan<br>FF | Yaeyama FF | Orii's FF | Daito FF | Philippines |
| --- | --- | --- | --- | --- | --- |
| Formosan FF |  | 0.000<sup>NS</sup> | 0.096* | 0.166* | 0.407* |
| Yaeyama FF | 0.011<sup>NS</sup> |  | 0.079* | 0.158* | 0.361* |
| Orii's FF | 0.057* | 0.036* |  | 0.247* | 0.519* |
| Daito FF | 0.132* | 0.139* | 0.159* |  | 0.439* |

<sup>NS</sup>: nonsignificant
\*: significant at $P < 0.05$

We found a significant positive correlation between pairwise genetic and geographical distances based on Mantel tests (*r*^2^ = 0.115, P < 0.05, Figure 2), indicating that genetic differentiation in *P. dasymallus* across the Ryukyu, Taiwanese and Philippine islands fits an isolation-by-distance model. Pairwise genetic distances also showed that the TW1 Formosan flying fox has a close relationship with Yaeyama populations from Iriomote-jima and Ishigaki-jima (0.082and 0.016, respectively), with pairwise distances that are lower than those between TW1 and both the TW2 and TW3 populations from the same subspecies (0.252 and 0.143, respectively).

**Figure 2.**
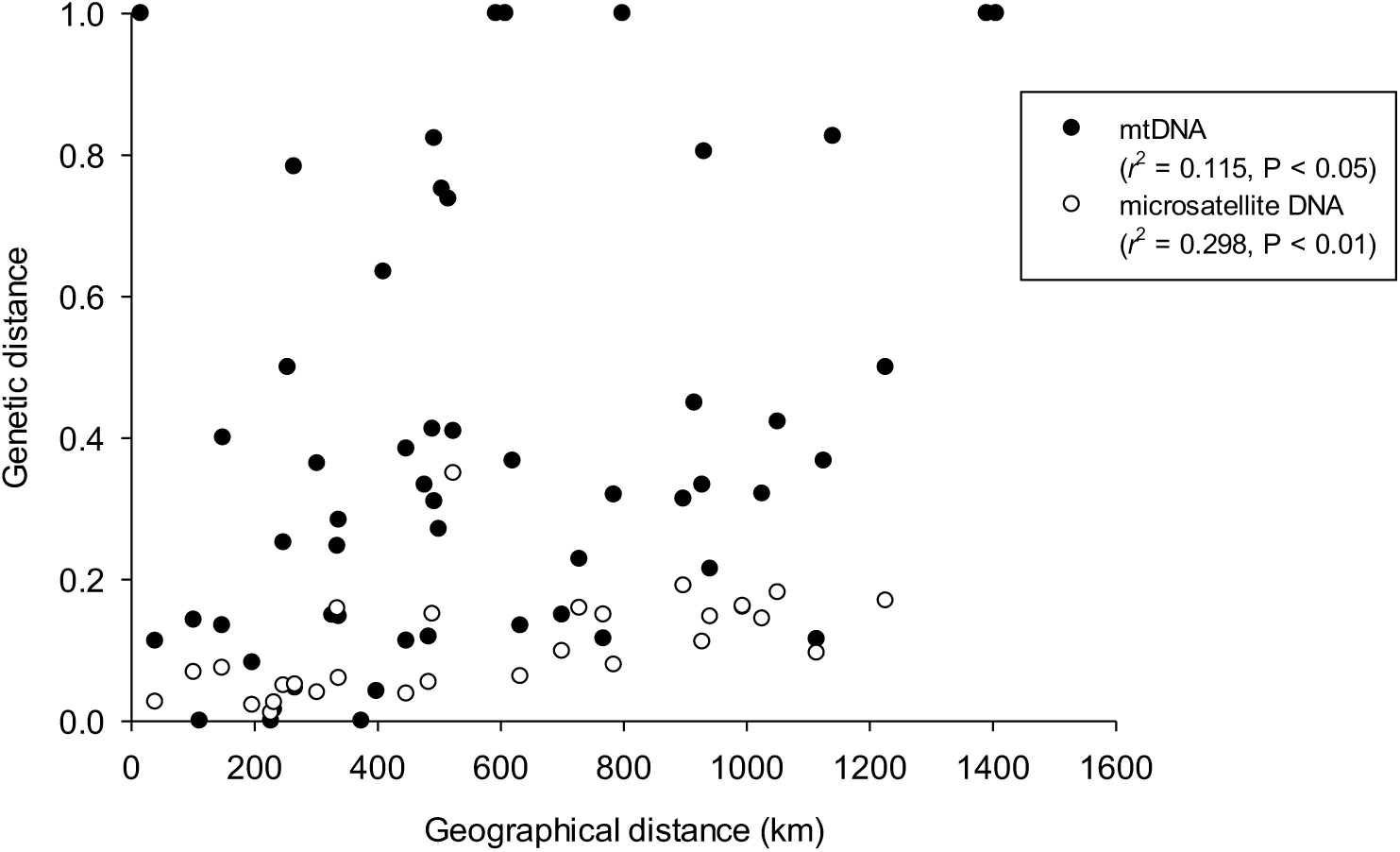
Relationships between genetic and geographical distances in *Pteropus dasymallus* in pairwise comparisons among islands.

The haplotype network indicated that one of the Philippine samples (MJV451) from Sabtang Island showed a relatively deeper genetic divergence with respect to the other samples. On the other hand, the haplotypes presented by Japanese or Taiwanese samples were genetically close to each other (Figure 3). Of these, two haplotypes were the most common, shared by 11and 10 individuals mainly found in Okinawa-jima (Orii’s flying fox) and Gueishan Island (TW1 Formosan flying fox), respectively.

**Figure 3.**
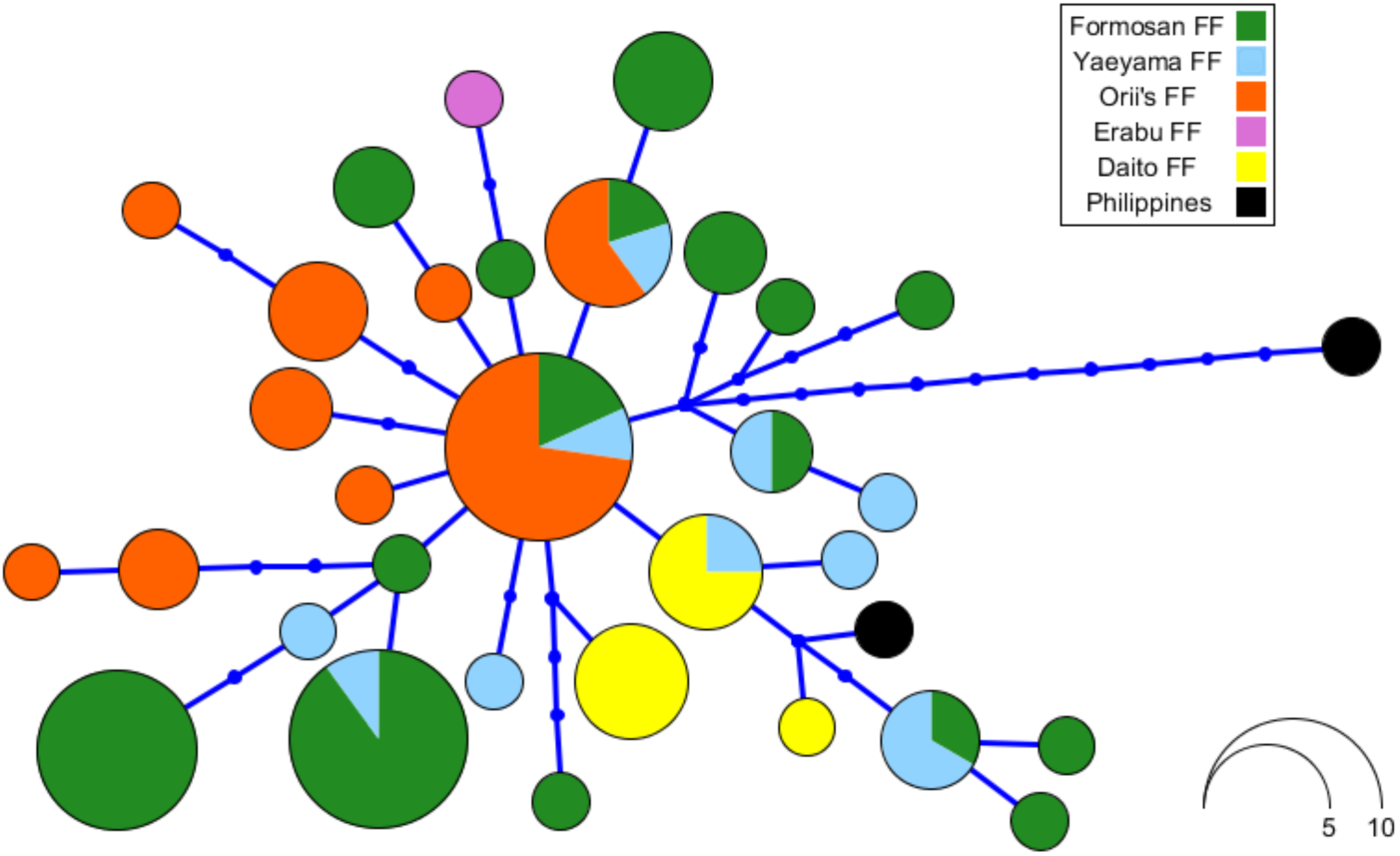
Haplotype network for *Pteropus dasymallus*. Each color represents a subspecies or population. The size of each circle is proportional to haplotype frequency. Each line segment and small dot represent a single mutational step and an inferred intermediate haplotype, respectively.

### Microsatellite analysis

Genotype analysis based on 26 polymorphic microsatellite loci from 76 *P. dasymallus* samples of five subspecies revealed a moderate degree of polymorphism across subspecies. The number of alleles at each locus was 5.27 ± 2.29, ranging from 2 to 10 across all the samples. The mean H_O_ and H_E_ values were 0.536 and 0.544, respectively. The highest diversity was recorded in the Formosan or Yaeyama flying fox. In contrast, and in line with the mtDNA data, the Daito flying fox harboured the lowest diversity (Table 1). The F_IS_ values were all not significant, implying no major deviations from HWE. Exact tests showed that four locus-population combinations deviated from HWE; however, there was no consistent pattern according to either subspecies or locus. No loci pair was detected in linkage disequilibrium. Finally, average pairwise relatedness was significant in three analyzed population except the Yaeyama. Daito flying fox showed a particularly high value of 0.147.

The AMOVA based on microsatellite data showed significant genetic differentiation among subspecies. The F_ST_ value was 0.069 (P < 0.001, Table 2). The Formosan and Yaeyama flying foxes were the only pair without significant differentiation. A pattern of isolation by distance was also shown here (*r*^2^ = 0.298, P < 0.01, Figure 2).

An analysis of genetic structure using STRUCTURE revealed clear substructure among geographical locations and subspecies identities (Figure 4). The most likely number of genetic clusters was four (*K*= 4) as inferred using the Evanno method based on the highest Δ*K* and mean likelihood value without an increase in variance. All or nearly all of the Daito and Orii’s flying fox samples, respectively, were assigned unambiguously to their own clusters, with the exception of one individual of Orii’s flying fox. A number of individuals also showed evidence of partial inferred ancestry. Formosan flying foxes from TW2 (from Lyudao) and TW3 (from Taiwan’s main island), and Erabu flying foxes, were assigned to a different genetic cluster. On the other hand, Formosan flying foxes from TW1 (from Gueishan Island) and Yaeyama flying foxes showed a greater admixture across different genetic clusters with full or partial membership. For *K* = 3, the Erabu and Orii’s flying foxes were grouped together. The Formosan and Yaeyama flying foxes showed an admixture of different genetic clusters. For *K* = 5, the Erabu flying fox also showed an admixture of membership.

**Figure 4.**
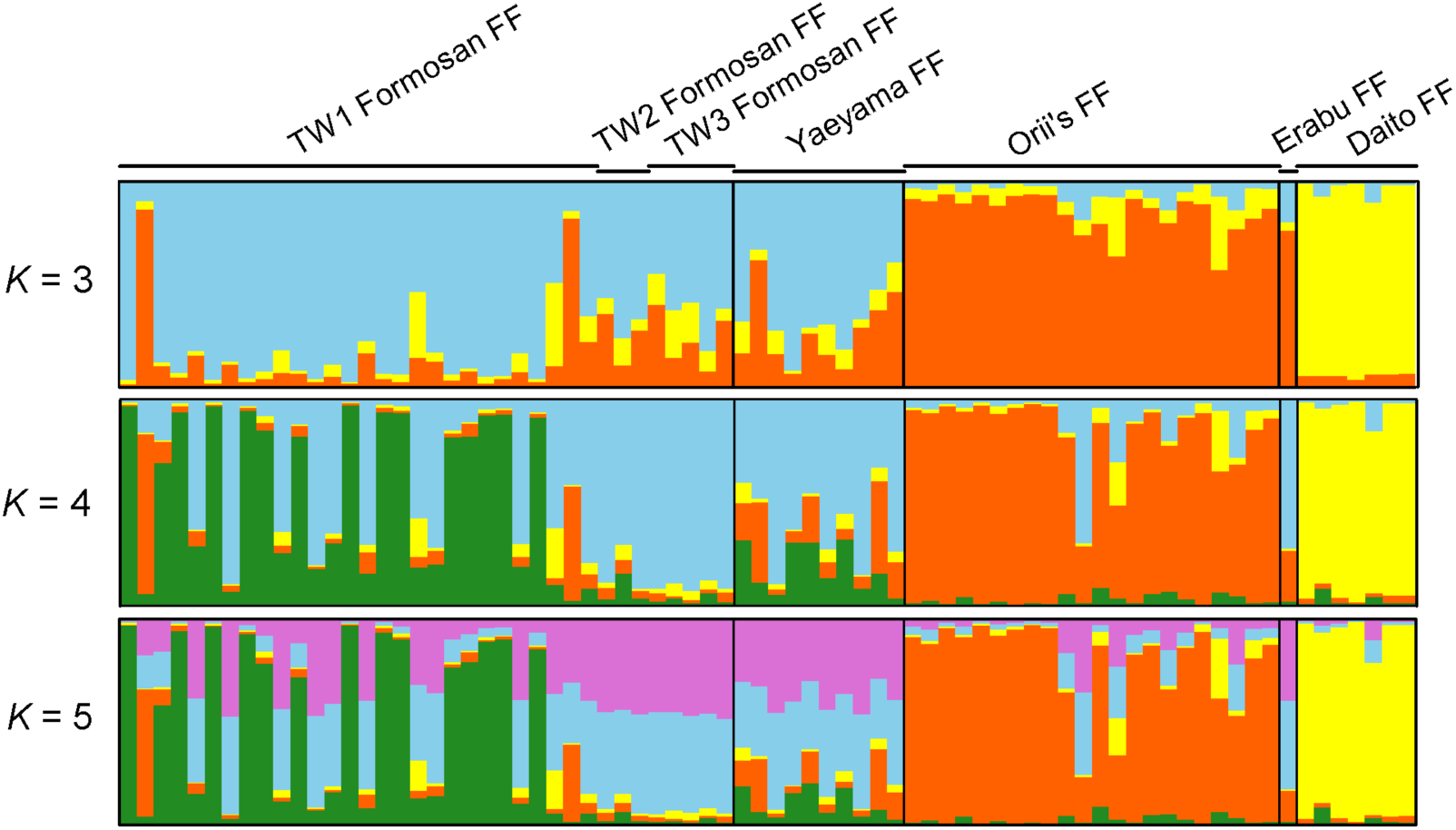
Genetic structure of 76 *Pteropus dasymallus* individuals from five subspecies based on a Bayesian STRUCTURE analysis. Each vertical bar represents one individual. Each color represents a genetic cluster. The length of each color in a certain vertical bar is proportional to the probability of assigning the individual to the corresponding cluster. The value of *K* indicates the possible number of clusters. The most likely number of clusters with the highest value of Δ*K* is four. The vertical black lines separate subspecies in different island groups. The Y-axis represents proportion, with a range between 0 and 1. Abbreviation: FF, flying fox.

## Discussion

We examined genetic diversity and structure among the five recognized subspecies of *P. dasymallus* from the Ryukyu Archipelago of Japan to Taiwan, and also included published data from two individuals sampled from a poorly known population from Batanes, Philippines. Our analyses based on mtDNA control region sequences and 26 microsatellite markers revealed significant genetic differentiation among island groups, broadly supporting the subspecies identities based on geographical locations (Figure 4).

Differences in the patterns of differentiation recovered by the two types of markers provide insights into the history of connectivity of these island populations. Notably, while we detected no deep divergence among any of the individuals from the five subspecies, the haplotypes showed evidence of only weak sorting with respect to island (Figure 3). Overall this pattern points to gene flow in the past, either through recurrent gene flow or colonization and admixture, alongside evidence of isolation and genetic drift in some cases, likely reflecting small population sizes. The central position of orange haplotype, which was most abundant in the Orii’s flying fox, suggests that this taxon might have served as a source of other populations across the Ryukyu and Taiwanese islands. In this scenario, the other subspecies populations were founded by colonization events, eastward to the Daito Islands and westward to Taiwan. While we were only able to examine two sequences from the Philippine population, these showed evidence of high levels of divergence with respect to each other, with the bat sampled from Sabtang Island also showing clear separation from all other samples. Further study, including sampling of bats from the Babuyan Islands, is needed to assess the likely causes of this apparent deep structure.

Our microsatellite analyses revealed a cline in genetic diversity from the highest in the Formosan and Yaeyama flying foxes, to the lowest in the Daito flying fox (Table 1). Of all the taxa, the Daito flying fox was seen to form a separate cluster with no admixture across different values of K. The genetic distinctiveness of the Daito flying fox can be explained by the comparatively large geographical distance between the remote easternmost Daito Islands and the other Ryukyu islands (approximately 360 km east off of Okinawa-jima) coupled with the absence of islands that could serve as stepping-stones for dispersers. The Daito Islands are uplifted coral islands that lie on the Philippine Sea Plate and are thought to have emerged approximately 1.2 to 1.6 million years ago in the mid-Pleistocene (Shiroma et al. 2015; Knez et al. 2017). Consequently, unlike the other continental islands, the oceanic Daito Islands have never been connected to a land mass by a land bridge during a glacial period. Instead, the Ryukyu Trench (Figure 1) – a deep, broad water body that separates the Daito Islands from the Eurasia Plate – has served as a significant geographical barrier to gene flow. We conclude that the low genetic diversity and high differentiation from other subspecies suggest the Daito subspecies arose from a historical event involving long-distance oceanic dispersal and has since experienced geographical and reproductive isolation. Similar differentiation and restricted gene flow between the Daito Islands and other Ryukyu islands lying on the Eurasia Plate has been reported for the elegant scops owl (*Otus elegans*) (Hsu 2005).

In addition to the Daito flying fox, we also found differentiation among other populations across the Ryukyu Archipelago based on multilocus genotypes, notably between the Erabu, Orii’s, and the Yaeyama flying foxes. In the case of Erabu, the inclusion of just one sample strongly limits our interpretations about this population. In contrast, the results for Orii’s and Yaeyama were more surprising, especially in light of the mtDNA data. Such subdivisions based on ncDNA are strongly concordant with the position of deep-sea channels, including the Tokara and Kerama Gaps, that separate the Northern, Central and Southern Ryukyus. Thus differentiation among these island groups, appears to have been driven by their long-term isolation from each other, a consequence of the fact that they were not connected by land-bridges during the Last Glacial Maximum.

In spite of the strong genetic subdivisions detected, our results also showed a significant positive correlation between genetic and geographical distance that is consistent with a pattern of isolation by distance across Taiwan and the Ryukyu Islands. Isolation by distance is typically considered to be a consequence of migration-drift equilibrium, whereby recurrent gene flow follows a stepping-stone pattern, and is thus more likely to occur among neighboring populations (Kimura & Weiss 1964). Nevertheless, trends of isolation by distance can also be generated through a colonization process, in which the contribution of genetic drift outweighs that of gene flow. Finally, drift and thus isolation by distance might also be more easily detected at larger spatial scales due to the higher probability that barriers will occur over greater distances (see Bossart & Prowell 1998). Indeed, in our study system, it is notable that island groups characterised by stronger genetic differentiation were also more likely to occur on opposite sides of deep-sea trenches (e.g., Hutchison & Templeton 1999).

Previous studies have reported gene flow among flying fox populations over hundreds to thousands of kilometres, although these have tended to focus on movements over land or along coast lines. In our study, similar genetic profiles of populations from Taiwan and Yaeyama support genetic mixing via movements across water, coupled with the formation of a land-bridge in the LGM. Indeed, these two subspecies are geographically closest and the least genetically structured. On the other hand, strong differentiation among island populations of flying foxes separated by 200-300 km, such as between Orii’s and Yaeyama, suggests that this distance represents an upper limit for recurrent gene flow in these bats. Nonetheless, despite overall clear genetic differentiation based on microsatellites, our structure-based clustering analyses did reveal a small number of putative migrants. In particular, two individuals recorded in Taiwan (TW1) and one in Yaeyama, appear to be individuals of the Orii’s flying fox.

A surprising result of this study was the high recorded genetic diversity in the Formosan flying fox from Gueishan Island (TW1). The inhabitants living on Gueishan Island before 1977, when the island was designated as a military control area, claimed that no flying fox had been seen on the island (Wu 2010), and thus this population is considered to be newly established via oceanic dispersal. Although this population appears to show a closer relationship with the Yaeyama flying fox from Iriomote-jima than with the other populations on Taiwan (TW2 and TW3), its high diversity likely stems from genetic admixture involving several different genetic clusters (e.g., Comas et al. 2004). Indeed, our results indicate that the TW1 population likely has multiple ancestral origins with putative founders from Yaeyama, TW2 (from Lyudao), TW3 (from the main island of Taiwan) and/or the Philippine population.

A combination of one or more explanations could account for the genetic diversity found on Gueishan Island. First, flying foxes might have arrived on Gueishan Island as a result of strong winds associated with seasonal typhoons or the winter northeast monsoon. Second, individual bats may have actively dispersed in search of resources. A third scenario is that active dispersal was driven by population expansion of the Yaeyama flying fox population. Yonaguni-jima of Japan, the westernmost margin of the distribution of the Yaeyama flying fox (Kinjo & Nakamoto 2009), is 107 km from Gueishan Island. According to Nakamoto et al. (2011a), Yaeyama flying foxes have been presumed to be dispersing eastward across the sea to a new insular habitat approximately 50 km away (from Tarama-jima to Miyako-jima). The flying fox population on Yonaguni-jima or the neighboring islands may also expand westward to Gueishan Island with the help of winds, forming a widely distributed and diverse population.

## Conclusions

Our findings from mitochondrial and nuclear markers support the current division of subspecies of *P. dasymallus* from the Ryukyu Archipelago and Taiwan. Genetic subdivisions among some island groups appear to reflect a lack of long-distance movements across water, coupled with the presence of deep-sea channels that prevented the formation of land-bridges during the LGM. We also find evidence that the recent colonization of Taiwan has involved founders from several distinct clusters. Taken together, we conclude that highly isolated and genetically distinct populations, such as Daito, should be treated as separate management units. On the other hand, bats from adjacent islands that show strong evidence of recent and frequent gene flow can be managed as a single population. The comparatively higher level of divergence between the Philippine sample from Sabtang Island and all the other sampled bats highlights the importance of future work to establish the status of this population. More generally, our results indicate that the evolutionary and ecological forces shaping the pattern of the genetic structure in *P. dasymallus* are dynamic and ongoing. As a taxon that ranges from the temperate northern Ryukyu Archipelago and subtropical Taiwan to the tropical northern Philippine islands, this species may serve as an excellent model for studying the processes driving island biogeography. Future studies that combine new sequencing technologies with more extensive sampling of the Philippine populations are expected to improve the current understanding of the phylogeography of *P. dasymallus* among islands.

## Acknowledgements

We are grateful to the Okinawa Zoo and Museum Foundation, Hirakawa Zoological Park, University of Ryukyus, Japan, and Taipei Zoo, Taiwan, for granting access to the collections. We thank Han-Chun Lee, Hui-Wen Wu, Ching-Lung Lin, Ching-Feng Lin, and many assistants for their hard work in the field. We also thank Dr. Si-Min Lin for valuable suggestions on the manuscript and Dr. Tetsuo Denda for laboratory support. This project was funded by the Forestry Bureau, Council of Agriculture (107-9.1-SB-17(1), 108-9.1-SB-30) and Ministry of Science and Technology (MOST 107-2621-B-305-001), Taiwan.

---

**Appendix 1.** Sample size and localities of origins of *Pteropus dasymallus* used in this study.

| Population name | # of samples | Localities of origins |
| --- | --- | --- |
| *P. d. formosus*<br>(Formosan flying fox) |  |  |
| TW1 | 28 | Gueishan Island |
| TW2 | 3 | Lyudao |
| TW3 | 5 | Taiwan's main island |
| *P. d. yayeyamae*<br>(Yaeyama flying fox) | 3 | Iriomote-jima |
|  | 7 | Ishigaki-jima |
|  | 1<sup>a</sup> | Irabu-jima |
| *P. d. inopinatus*<br>(Orii's flying fox) | 22 | Okinawa-jima |
| *P. d. dasymallus*<br>(Erabu flying fox) | 1 | Kuchinoerabu-jima |
| *P. d. daitoensis*<br>(Daito flying fox) | 7 | Minamidaito-jima |
| Philippine population | 1<sup>b</sup> | Batan Island |
|  | 1<sup>b</sup> | Sabtang Island |
| Total | 80 |  |

<sup>a</sup>: sequence acquired from Genbank with an accession numbers of NC\_002612.1 (Nikaido et al. 2000)
<sup>b</sup>: sequences acquired from Dryad with catalog numbers of MJV451 and MJV458 (Tsang et al. 2019)

## Supporting Information

**S1.**
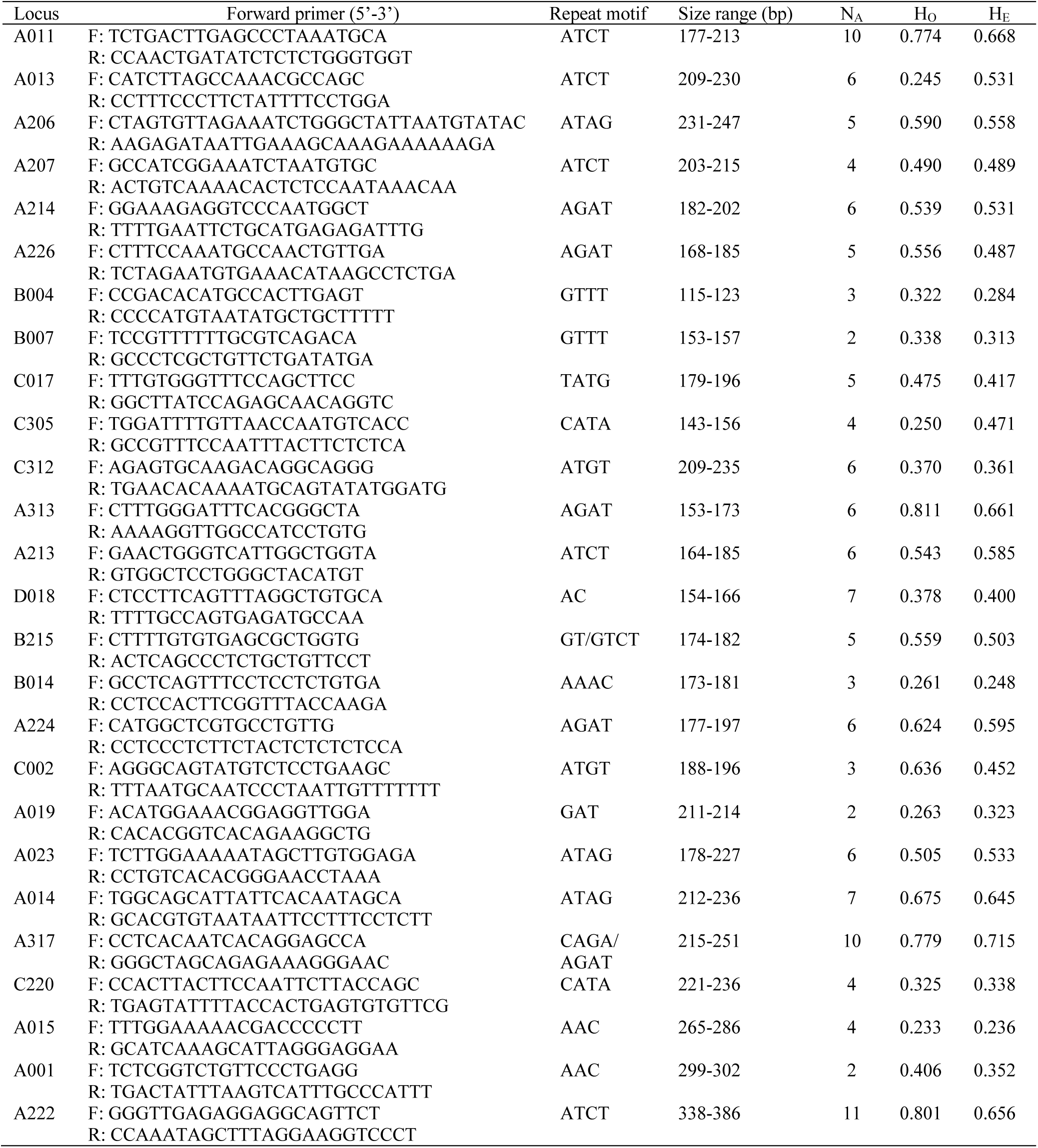
Twenty-six polymorphic microsatellite loci for *Pteropus dasymallus* used in this study. N_A_: number of alleles; H_O_: observed heterozygosity; H_E_: expected heterozygosity.

